# Hepatitis B virus envelope proteins can serve as therapeutic targets embedded in the host cell plasma membrane

**DOI:** 10.1101/2020.12.21.423802

**Authors:** Lili Zhao, Fuwang Chen, Oliver Quitt, Marvin Festag, Marc Ringelhan, Karin Wisskirchen, Julia Hasreiter, Luidmila Yakovleva, Camille Sureau, Felix Bohne, Michaela Aichler, Volker Bruss, Maxim Shevtsov, Maarten van de Klundert, Frank Momburg, Britta S. Möhl, Ulrike Protzer

**Author notes:** Corresponding author: Prof. Ulrike Protzer, MD, Institute of Virology, Trogerstr. 30, 81675 Munich, Germany. These authors contributed equally to this work (Oliver Quitt, Marvin Festag).

## Abstract

Hepatitis B virus (HBV) infection is a major health threat causing 880,000 deaths each year. Available therapies control viral replication, but do not cure HBV leaving patients at risk to develop hepatocellular carcinoma. Here we show that HBV envelope proteins (HBs) - besides their integration into endosomal membranes - become embedded in the plasma membrane where they can be targeted by redirected T-cells. HBs was detected on the surface of HBV-infected cells, in livers of mice replicating HBV and in HBV-induced hepatocellular carcinoma. Staining with HBs-specific recombinant antibody MoMab recognizing a confirmational epitope indicated that membrane-associated HBs remains correctly folded in HBV-replicating cells in cell culture and in livers of HBV-transgenic mice *in vivo*. MoMab coated onto superparamagnetic iron oxide nanoparticles allowed to detect membrane-associated HBs after HBV infection by electron microscopy in distinct stretched of the hepatocyte plasma membrane. Last not least we demonstrate that HBs located to the cell surface allows therapeutic targeting of HBV-positive cells by T-cells either engrafted with a chimeric antigen receptor or redirected by bispecific, T-cell engager antibodies.

## Introduction

The human hepatitis B virus (HBV) is the prototypical member of the *Hepadnaviridae* family. The virion consists of an icosahedral capsid containing the viral DNA genome. The capsid is enveloped by a cell-derived lipid bilayer carrying the three HBV envelope proteins. Expression of the three isoforms is regulated via transcriptional and translational initiation. The N-terminally extended variants of the small envelope protein (S), referred to as middle (M) and large (L) envelope proteins, share the S protein domain and collectively are referred to as hepatitis B surface (HBs) proteins. Beside infectious virions, an excess of subviral particles (SVPs) is produced and secreted in 1,000-100,000-fold excess over virions (Blumberg, 1977). SVP contain densely packed, membrane embedded HBs but do not contain viral DNA. SVP are released in the form of spheres or filaments from infected cells or cells that have integrated HBV-DNA and can be detected as HBsAg in patient blood. Presumably, HBsAg secreted in excess over virions benefits viral persistence by limiting the development of an effective adaptive immune response.

Worldwide, more than 250 million individuals are chronically infected with HBV. Chronically infected individuals have an increased risk of developing liver fibrosis, cirrhosis and hepatocellular carcinoma (HCC), leading to an estimated 880,000 HBV infection-related deaths per year (WHO, 2017). Available antiviral therapy with nucleos(t)ide analogues suppresses viral replication, but does not cure the infection and has limited impact on the development of hepatocellular carcinoma (Chen, Wang et al., 2017, Debarry, Cornberg et al., 2017, Lobaina & Michel, 2017, WHO, 2017). Therapy with (pegylated) interferon (IFN) alpha is limited by severe side effects. Therapy-induced or spontaneous loss of detectable serum HBsAg, regarded as functional cure, is rare, but almost completely leverages the odds of developing end-stage liver disease. Therefore, it is anticipated that a therapy that cures HBV would greatly reduce HBV-infection related morbidity and mortality.

The S-domain is common for the three envelope proteins and carries the major hydrophilic determinant including the ‘a’ determinant, an extracellular antigenic loop, which defines the antigenic properties of HBsAg, i.e. SVPs and virions, of different HBV genotypes (Norder, Courouce et al., 2004). Different antibody binding properties in the “a” determinant of different HBV genotypes lead to the definition of HBV serotypes. The extraordinary conformation of this antigenic determinant is composed of three loops formed by numerous disulfide bonds (Bhatnagar, Papas et al., 1982, Brown, Howard et al., 1984, Dreesman, Sanchez et al., 1982, Mangold, Unckell et al., 1995, Qiu, Schroeder et al., 1996). The L protein carries an N-terminal myristoylation as infectivity determinant and the receptor binding regions for the interaction with heparan sulphate proteoglycans (HSPGs) and the bona fide HBV receptor, the bile acid transporter sodium-taurocholate co-transporting peptide (NTCP) (Leistner, Gruen-Bernhard et al., 2008, Ni, Lempp et al., 2014, Schulze, Gripon et al., 2007, Sureau & Salisse, 2013, Yan, Zhong et al., 2012).

Infectious virions and filamentous SVP are assembled by budding into the endoplasmic reticulum, translocate to multivesicular bodies (MVB), and are released via the endosomal sorting pathway (Hoffmann, Boehm et al., 2013, Jiang, Himmelsbach et al., 2015, Lambert, Doring et al., 2007, Stieler & Prange, 2014, Watanabe, Sorensen et al., 2007). Immunohistochemistry (IHC) analysis of liver tissue sections has revealed that in chronically HBV infected patients, a significant portion of the HBs produced in infected hepatocytes may localise to the cell membrane (Busachi, Ray et al., 1978, Chu & Liaw, 1995, Safaie, Poongkunran et al., 2016a, Safaie, Poongkunran et al., 2016b). This suggests that with assembly and release of HBV virions and filaments from MVBs a proportion of the surface antigens become embedded into the MVB membrane, and subsequently, upon fission between the MVB and the cell membrane, ends up in and on the cell membrane.

In the current study, we aim at characterizing the intracellular localization of the HBV envelope proteins in HBV-replicating and -infected hepatoma cells as well as in liver specimens using different HBs-specific antibodies. We found that HBs localises to the plasma membrane of HBV replicating cells in its natural conformation. We developed and characterised the conformation-specific, humanised anti-HBs antibody MoMab, and showed that MoMab can detect membrane-associated HBs on HBV infected hepatocytes by immunofluorescence staining and transmission electron microscopy (TEM), and in liver tissue of HBV-transgenic mice. Furthermore, we demonstrate that membrane-localised HBs can be targeted by redirected T-cells thus providing an interesting therapeutic target.

## Results

### HBs localises to the plasma membrane *in vivo*

As previous studies indicated that HBs maybe localisced on the membrane of hepatocytes (Hoffmann et al., 2013, Jiang et al., 2015, Lambert et al., 2007) and membranous staining of HBs on hepatocytes strongly correlated with active viral replication (Jiang et al., 2015), we evaluated tissue section obtained from livers of HBV-transgenic mice constitutively replicating HBV from integrated, 1.3-fold overlength HBV genome (Guidotti, Matzke et al., 1995) and from HBV-infected patients who have developed hepatocellular carcinoma. Immunohistochemistry (IHC) staining using HBs-specific antibody 70-HG15 revealed that the extent of HBs staining in hepatocytes shows large interindividual variation but also varies from cell to cell (Fig 1, EV1). Accumulation of HBs on the cell membrane was observed in cells with both, high- and low amounts of intracellular HBs in transgenic mice in which each hepatocyte contains the same transcription template (Fig 1A). In liver biopsies from chronic hepatitis B patients, HBs staining was highly variable from cell to cell distribution and showed a mixed pattern (Fig 1B, EV1), but staining indicative for a membranous localization was found in the presence or absence of concomitant cytoplasmic staining. Taken together, our IHC results confirmed the speckled membranous-localization of HBs that has been described before in liver tissue.

**Figure 1.**
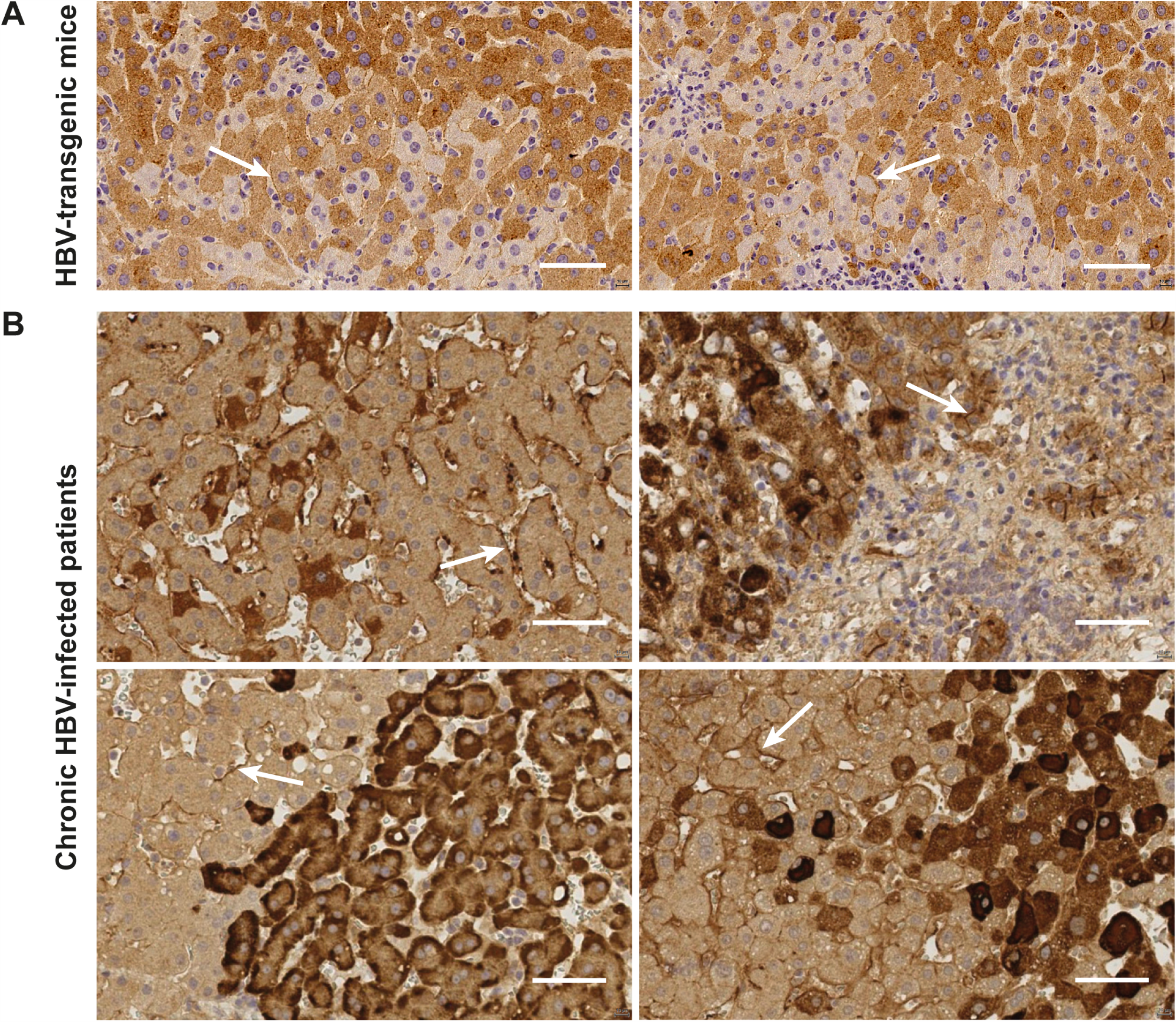
Intrahepatic HBs distribution and localisation in liver sections. Immunohistochemistry staining for HBs to determine distribution in liver tissue of (A) HBV-transgenic mice and (B) hepatocellular carcinoma resected from HBV-infected humans. HBs was stained using the HBs-specific antibody 70-HG15. Membrane-associated HBs is indicated by white arrows. Scale bars represent 50 μm.

### Membrane-associated HBs is derived from endogenously expressed protein

To determine whether membranous HBs is derived from endogenously expressed protein translocated to the plasma membrane, we stained HBs by immunofluorescence on HepG2.2.15 and HepAD38 cells, which constitutively replicate HBV. To prevent detection of intracellular HBs, cells were stained with different anti-HBs antibodies (HBVax, HB01, 5F9) prior to permeabilization. Co-staining with wheat germ agglutinin (WGA) revealed that HBs localises to the plasma membrane on HepG2.2.15 cells (Fig 2A), as well as HepAD38 cells (data not shown). Membranous HBs distribution was characterised by a distinct punctuate pattern (Fig 2A). When parental HepG2 cells were incubated with supernatant of HepG2.2.15 cells prior to staining with anti-HBs antibody 5F9 to allow binding of secreted SVP to the cell membrane, we were not able to stain HBs on the cell membrane (Fig 2B). This indicates that the membrane-associated HBs on HepG2.2.15 cells is derived from intracellularly expressed HBs.

**Figure 2.**
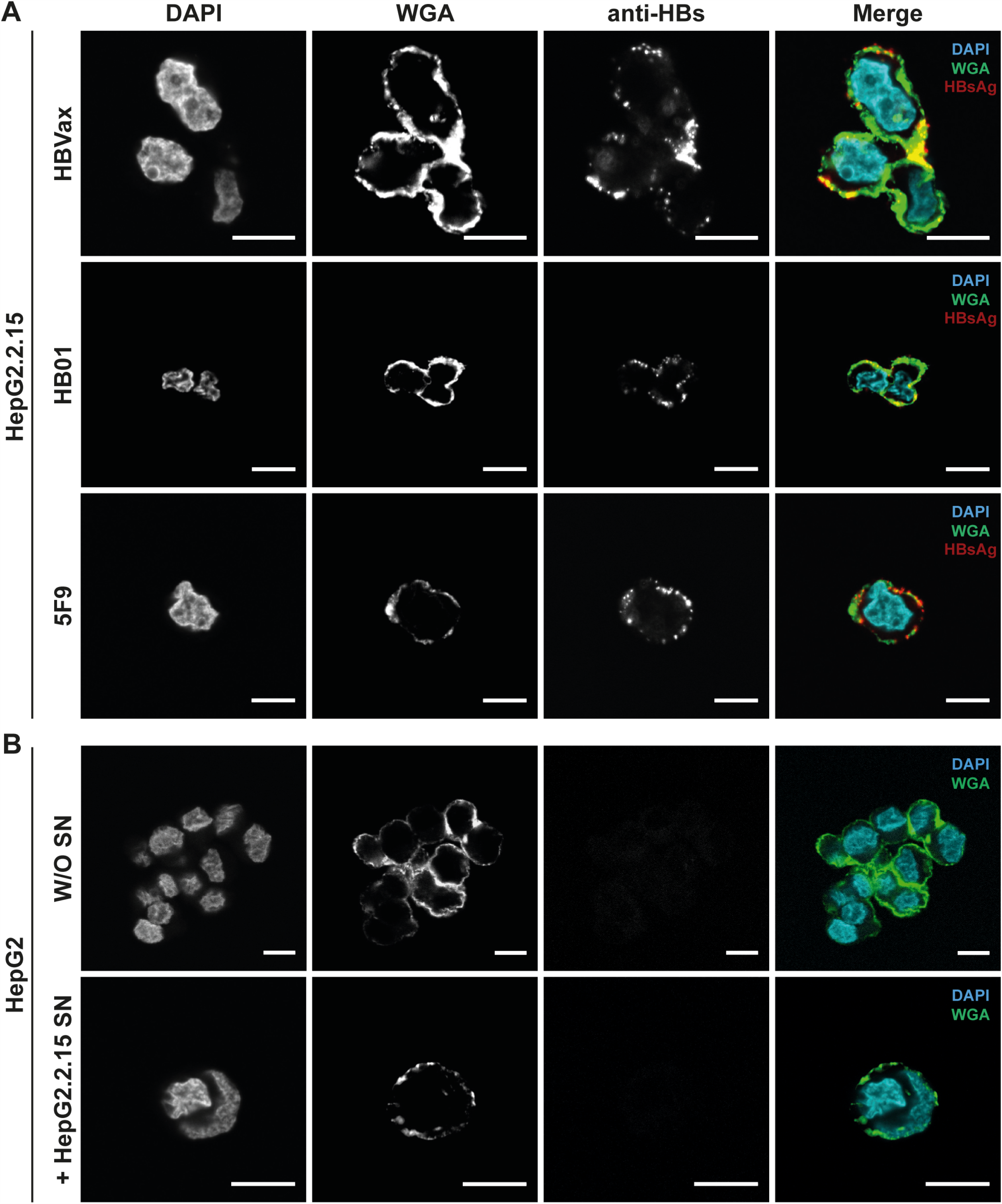
Membrane associated HBs is derived from endogenously expressed HBs. A HepG2.2.15 cells, which constitutively replicate HBV, were incubated with the indicated HBs specific antibodies before permeabilization. Confocal microscopic analysis shows that HBs (white, merged images on the right, red) co-localises with the cell membrane, which was stained with wheat germ agglutinin (WGA, green). B HepG2 cells were incubated with HepG2.2.15 supernatant (SN) prior to HBs staining with 5F9. Neither staining of HepG2 cell without (W/O SN) nor after incubation with HepG2.2.15 SN detected HBs on the cell membrane. Chromatin was co-stained by DAPI (cyan). Exemplary representative staining’s of five independent experiments are shown. Scale bars represent 10 μm.

Next, Huh7 cells were transfected with plasmid DNA expressing either the S or L HBV envelope protein, or all HBs isoforms combined (SML) (Fig 3A). Transfected cells were lysed and supernatants were collected at 24, 48, and 72 hours post-transfection. Protein expression and secretion was confirmed by Western blot analysis (Fig 3B). For sensitive detection of secreted HBsAg the Architect™ HBsAg QT assay was applied (Fig 3C). In line with observations by others, the L protein, when expressed in absence of the S protein, was not secreted (Fig 3B, 3C). We observed comparable glycosylation of the envelope proteins expressed from the different vectors. Expression levels of L were low when no S and M were co-expressed and L alone was not secreted (Fig 3B). Immunofluorescent staining of non-permeabilized Huh-7 cells transfected with the S-, L- and SML-expressing plasmid vectors revealed that in all cells HBs is associated with the plasma membrane in a distinct, punctuate pattern. Exemplary staining using the HBs-specific antibody HB01 is shown (Fig 3D). The observation that L, although it cannot be secreted in SVP on its own, locates to the plasma membrane confirms that HBs detected on the cell surfaces is derived from endogenously expressed HBs.

**Figure 3.**
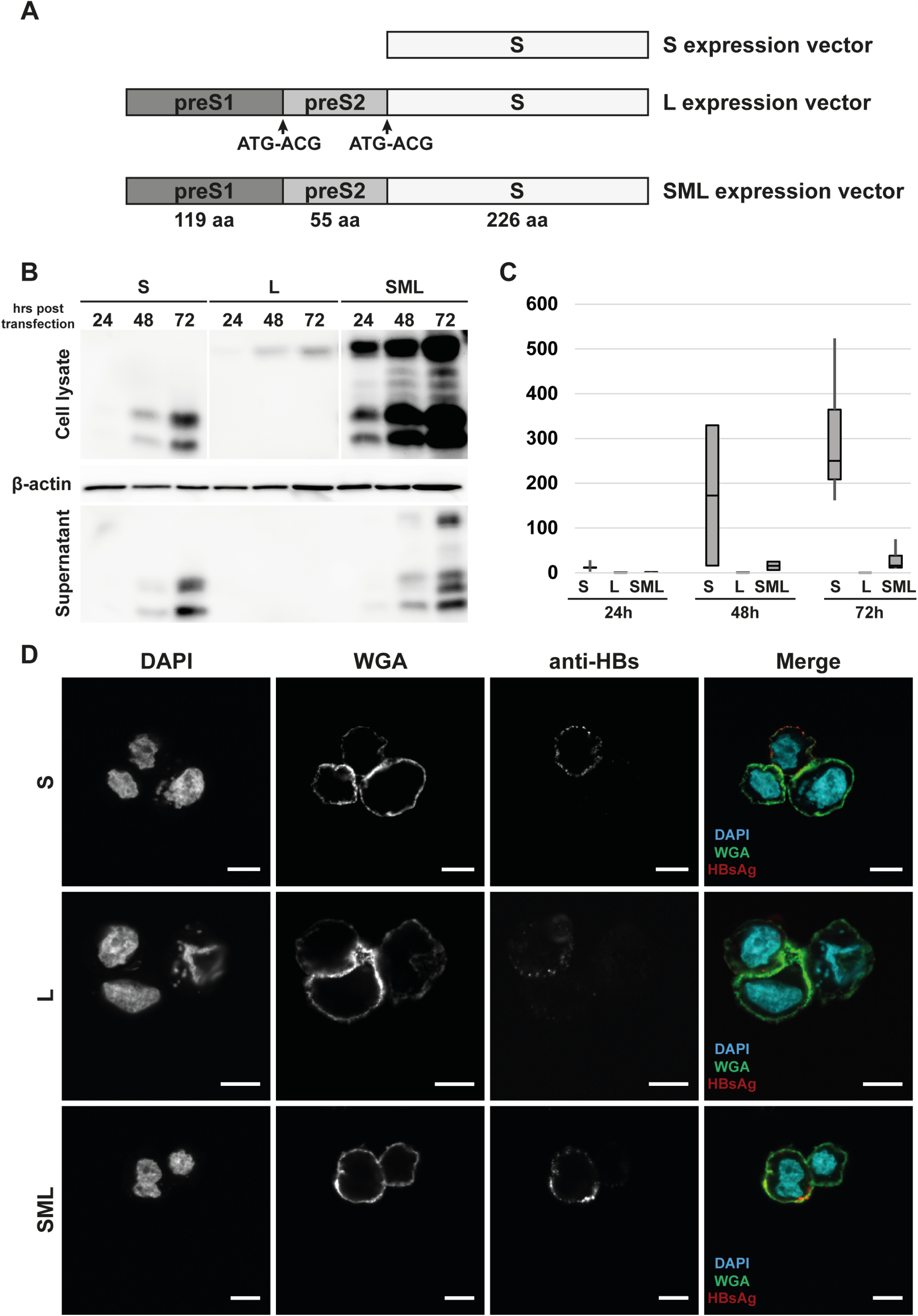
HBs are embedded in the cell membrane independent of SVP secretion. A Schematic overview of plasmids expressing only HBV S- or L-protein or all three envelope proteins (SML) of HBV genotype A. To express the L protein in absence of M and S proteins, the respective start codons were mutated (ATG-ACG, indicated by arrows). B Western blotting analysis of the cell lysates (upper panel) and supernatants (bottom panel) of transfected Huh7 cells using a monoclonal, S-specific antibody (HB01). S, M and L proteins were expressed in transfected Huh-7 cells in non-glycosylated and glycosylated forms at 24, 48 and 72 hours post transfection. Beta-actin was used as a loading control. C HBsAg was quantified in the supernatant of transfected Huh-7 cells using the Architect™ HBsAg QT assay. HBs could be detected in the supernatant of cells expressing the S or SML protein(s), but not the L protein alone. Graph represents the average and mean ± SD of three independent experiments. D Confocal microscopic analysis of non-permeabilized S-, L- or SML-transfected Huh7 cells confirmed the speckled membrane-localisation of HBs (white, merged images: red). The plasma membrane was co-stained using WGA staining (green), and chromatin by DAPI-staining (cyan). Exemplary representative staining’s of two independent experiments are shown. Scale bars represent 10 μm.

### Characterisation of the new humanised HBs-specific antibody MoMab

To characterise membrane-associated HBs in detail, we constructed a chimeric, human antibody designated as MoMab. MoMab is a homo-dimer of a human single chain antibody fragment (scFv C8) (Bohne, Chmielewski et al., 2008) directed against the S-domain of all three HBV envelope proteins that has been terminally fused to the constant region (CH2 and CH3) of human IgG1 (Fig 4A). The scFv C8 originates from a scFv library that was derived from peripheral blood lymphocytes of individuals vaccinated against hepatitis B and was selected because of its high affinity to HBs-expressing hepatoma cells. MoMab was expressed in HEK 293 cells and purified from cell culture supernatant by ion exchange chromatography. Coomassie blue staining of purified MoMab separated by polyacrylamide gel electrophoresis under reducing and non-reducing conditions indicated that the purified MoMab was free of other protein contaminations (Fig EV2A). Binding affinity of purified MoMab that contains two HBs-binding domains (Fig. EV2A) to immobilised HBsAg was determined by ELISA, revealing an affinity constant of 0.64 nM (Fig 4B).

**Figure 4.**
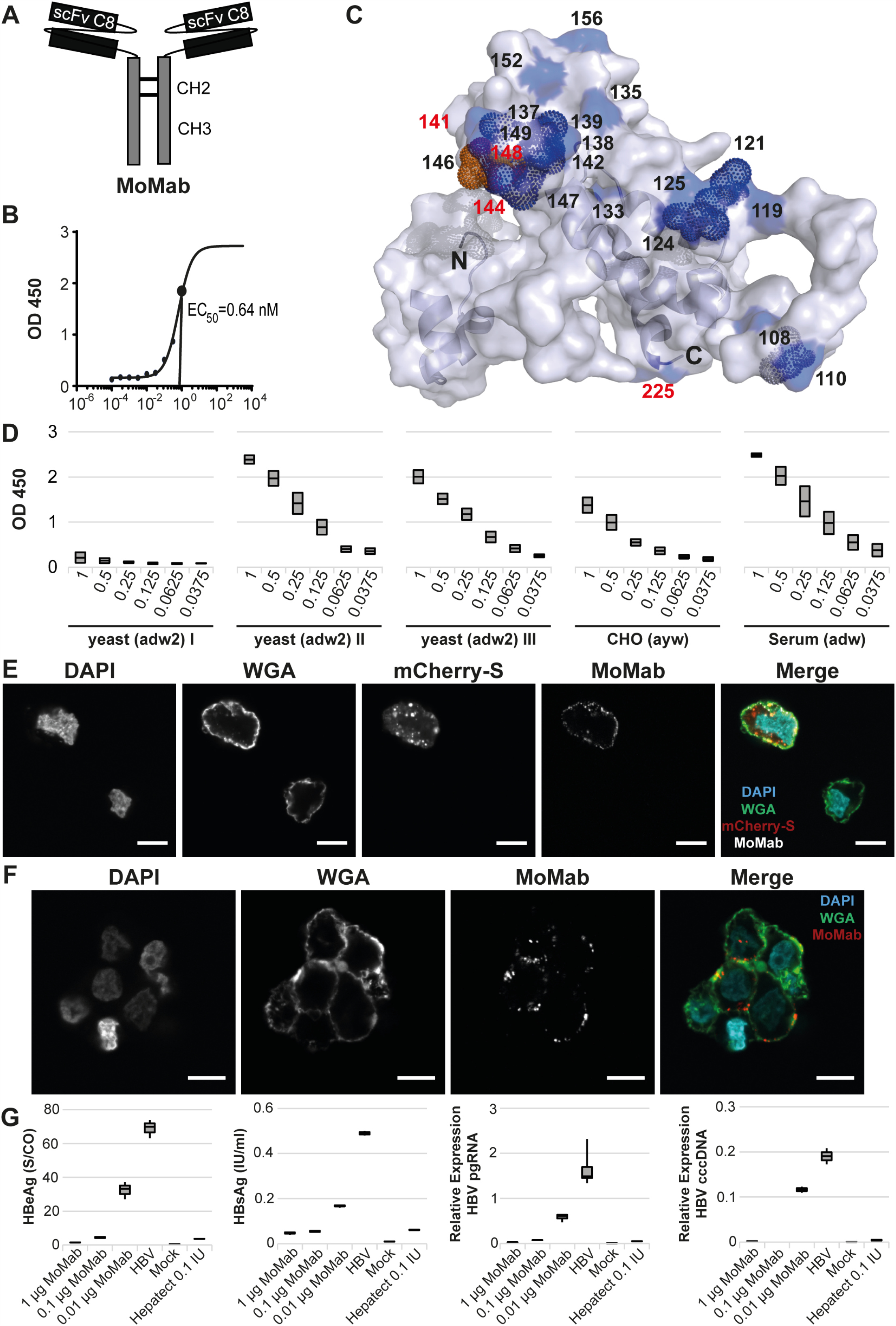
Generation and characterisation of a human, recombinant antibody for detection of HBs. A Schematic representation of the recombinant human antibody MoMab. A human single chain antibody fragment (scFv C8) was N-terminally fused to the constant region (CH2 and CH3) of human IgG1 heavy chain replacing VH and CH1 domains. B The avidity of MoMab for immobilised HBsAg purified from patients’ serum was determined by ELISA, revealing a 50% functional affinity constant (EC50) of 0.64 nM. C HBs residues that are involved in the interaction between MoMab and HBs were identified by an alanine scanning analysis (see Figure EV2B) and mapped to a theoretical model of the HBV S structure (pdb file modified from van Hemert et al., 2008). Amino acids of S that are part of the MoMab epitope are shown in blue. Numbers indicate amino acid positions, numbers in red indicate residues most critical for the interaction. Cystein mutations that disrupted the interaction between MoMab and HBs and the glycosylation site N146 (orange) are shown as dotted spheres. The N- and C-terminus are indicated with N and C, respectively. D HBsAg from different serotypes and divergent origins (yeast, CHO cells and serum) was immobilised on ELISA plates at different concentrations. The capacity of MoMab to interact with immobilised antigen was determined by ELISA. Graph represents the average and mean ± SD of two experiments performed in triplicate. E HepG2 cells were transfected with a plasmid expressing an mCherry-S fusion protein (red). F HepG2-NTCP cells were infected with HBV at MOI 500. Cells were stained with MoMab coupled to Alexa-Fluor® 647 fluorochrome (white) before fixation and permeabilization. The cell membrane was co-stained with WGA (green) and cell nuclei were stained with DAPI (Blue). Representative pictures obtained in two independent experiments are shown. Scale bars represent 10 μm. G For the neutralisation assay, purified HBV was incubated with different amounts of MoMab and applied to infect HepG2-NTCP K7 cells at MOI 100. HBeAg and HBsAg levels were determined in cell culture supernatant collected from day 4 to 8 post infection and are given as signal over cut-off (S/CO) and international units (IU) per ml, respectively. Cells were lysed 12 days post infection and intracellular HBV pregenomic RNA (pgRNA) and cccDNA were determined by qPCR. Relative expression levels after normalization to housekeeping gene β-actin are shown. Graphs represent the mean ± SD of triplicate infection experiments.

Next, we determined the epitope within the S-protein domain targeted by MoMab using an alanine-scanning approach. Point mutants of S within its “a” determinant (aa 101 to 225) were expressed in mammalian cells, immobilised and exploited to assess MoMab binding (Fig EV2B). Residues that, when mutated, disrupted the interaction between MoMab and HBs, were mapped on to the hypothetical structure of the S protein (van Hemert, Zaaijer et al., 2008), revealing that MoMab targets the antigenic loop of the ‘a’ determinant (Fig 4C, EV2B). The largest impacts on MoMab binding to HBs were monitored for the major immune-dominant residues such as the charged residues K141 and D144 as well as the non-charged, polar residue T148 within the antigenic loop (Fig 4C, S2B). Interestingly Y225, which is highly-conserved among all HBV genotypes except genotype F, was also an important residue determining the confirmational epitope recognised by MoMab (Fig 4C, EV2B). ELISA analysis using the antibody on immobilised HBsAg indicates that MoMab recognises HBsAg from different serotypes and divergent origins such as yeast, Chinese hamster ovary (CHO) cells and human serum (Fig 4D). However, MoMab did not recognise HBsAg (Fig 4D), which is known to have lost its native structure. All these results highlight that MoMab binding is highly confirmation dependent.

To show that the relatively low amounts of HBs ending up on the cell surface suffice to bind the MoMab, we stained HepG2 cells transfected with a plasmid expressing a mCherry-S fusion protein as well as HBV-infected HepG2-NTCP cells. As the cells were not permeabilized, the abundant HBs accumulating in ER membranes could not be targeted by the MoMab added. Purified MoMab specifically interacted with HBs on the surface of mCherry-expressing HepG2 cells but not with neighbouring HBs-negative cells (Fig 4E, EV2C) and with HBV-infected HepG2-NTCP cells (Fig 4F, EV2D). Staining intensity increased with the MOI used for infection of HepG2-NTCP cells (Fig EV3). To assess whether MoMab has neutralising activity, purified HBV was pre-incubated with different concentrations of MoMab (1µg, 0.1µg and 0.01 µg) before infection of HepG2-NTCP cells. Pre-incubation with MoMab dose dependently inhibited HBV infection as shown by a decrease of HBeAg and HBsAg secretion 8 days post infection, as well as intracellular HBV pregenomic RNA and cccDNA determined 12 days post infection (Fig 4G).

Taken together, these data show that the chimeric, recombinant human MoMab antibody detects a confirmational epitope within the “a” determinant of the S-domain of HBV envelope proteins of different serotypes with high avidity and is able to neutralise HBV infection. It sensitively detects HBs on the surface of HBs-expressing and HBV-infected cells.

### Detection of HBs by MoMab conjugated nanoparticles

To visualise membrane-associated HBs on the cell surface and exclude detection of HBV particles bound to the cell surface, MoMab was conjugated to dextran-coated superparamagnetic iron oxide nanoparticles (SPIONs) (Fig EV4A) allowing detection by transmission electron microscopy (EM). A ‘switch’ assay of MoMab-conjugated SPIONs co-incubated with soluble HBsAg demonstrated correct formation of antibody conjugates by an increase in the T2 relaxation time (Fig EV4B, C). MoMab-conjugated SPIONs were incubated with constitutively HBV-replicating cells (HepG2.2.15 and HepAD38), and HBV-infected HepG2-NTCP cells. EM analysis demonstrated that MoMab-SPIONs bind the plasma membrane of all HBV replicating cells (Fig 5). In accordance with the punctuate in immunofluorescence stainings we observed (Fig. 4E, F) HBs was concentrated at certain membrane areas. There was little indication for binding to viral or sub-viral particles. HBs visualised by MoMab-SPION binding was distributed at certain membrane stretches on cell protrusions where they tend to cluster.

**Figure 5.**
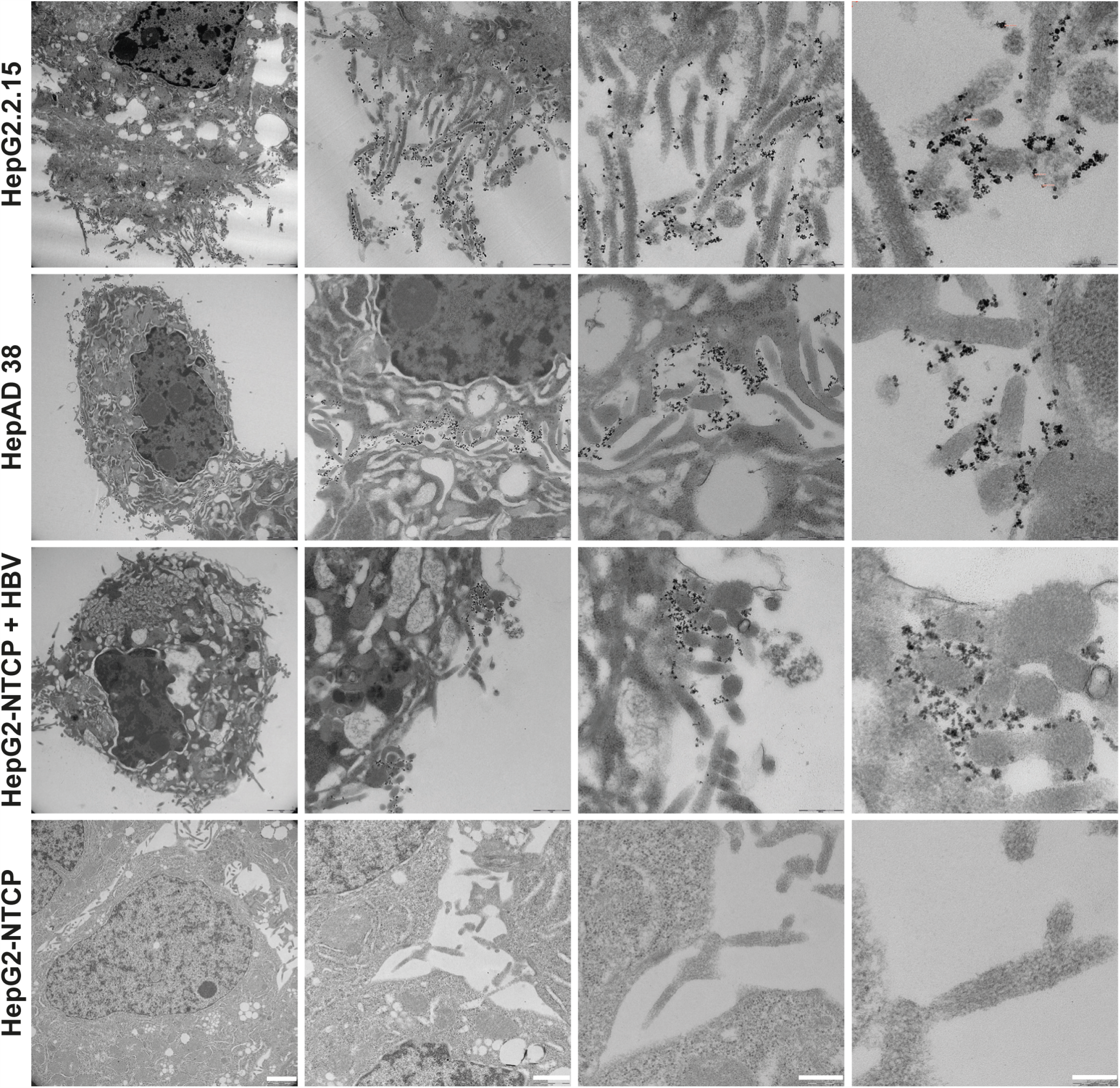
MoMab-coupled iron particles indicate localisation of HBs to distinct membrane stretches on the hepatocyte surface. MoMAb was coupled to SPIONs and incubated on HepG2.2.15 (top row) or HepAD38 cells (second row) constitutively replicating HBV, HBV infected HepG2-NTCP cells (third row) and non-infected control cells (fourth row). After fixation, transmission electron microscopy of ultrathin sections was performed. Cells visualised by electron microscopy at increasing magnification (from left to right) showed binding of MoMab-SPIONs localising to the membranes on protrusions of HBV replicating, but not of HBV-negative HepG2-NTCP cells. Scale bars (white) from the left to the right indicate 2 μm, 1 μm, 500 nm and 200 nm.

### HBs on the cell surface allows therapeutic targeting by CAR-T cells or T-cell engager antibodies

To assess whether the MoMab would target membrane-associated HBs *in vivo*, Alb-Psx transgenic mice that express HBV L protein under control of an albumin promoter in hepatocytes were intravenously injected with 100 µg MoMab. Because mainly L protein, but little S protein is expressed, these mice have low or even no detectable HBsAg levels in their blood. Four hours after MoMab injection, HBsAg level in serum and intrahepatic HBs was quantified. In one mouse with detectable HBsAg level, serum HBsAg decreased from 43.35 IU/ml to 8.95 IU/ml after MoMab injection (data not shown). In this mouse, MoMab was detected in the liver by staining with an anti-human antibody, indicating that MoMab targeted HBs expressed on hepatocytes *in vivo* (Fig 6A). MoMab did not target hepatocytes of wild-type C57BL/6 mice (Fig 6B).

**Figure 6.**
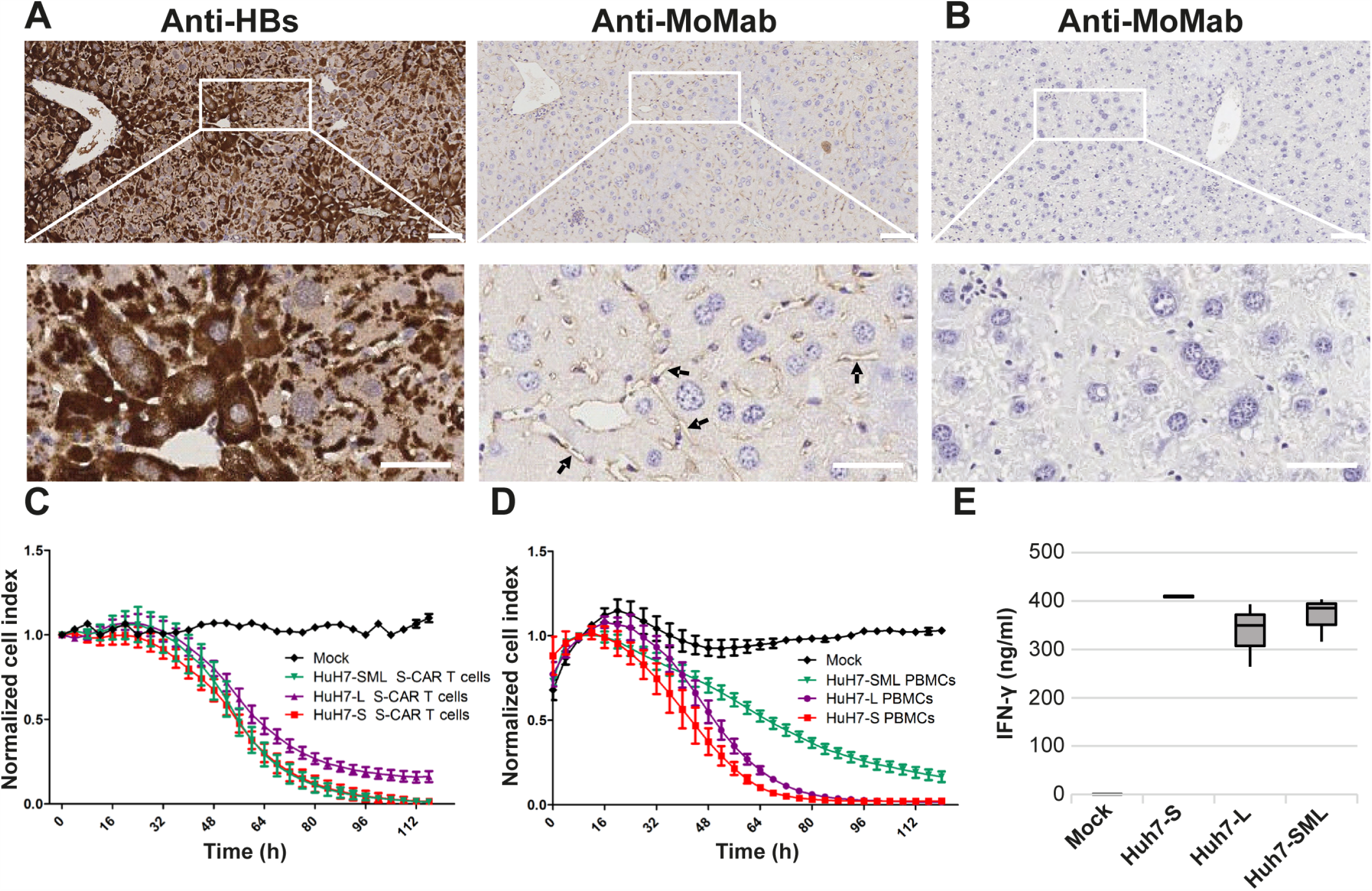
Targeting of membrane-localised HBs by antibodies and CAR-T cells. A 100μg purified MoMab were i.v. injected into four AlbPsx mice that expresses HBV L protein. After 4 hours, mice were sacrificed and livers collected. HBs was stained using anti-HBs antibody 70-HG15 or an anti-human antibody to detect the MoMab that had been injected. B MoMab-staining of liver sections of WT-mice served as negative control. Scale bars represent 50 μm. C, D Huh-7 cells were transfected with plasmid DNA to express only HBV S- or L-protein, or all three envelope proteins (SML). C Transfected cells were co-cultured with human T-cells engrafted with a scFv C8 based chimeric antigen receptor (S-CAR). Cell viability was followed by real time cytotoxicity analysis for 4 days using an xCelligence device. Normalised cell index indicated that the engrafted T-cells specifically killed all HBs-expressing Huh-7 cells. D Transfected HuH7 cells were co-cultured with human PBMCs in the presence of a bispecific T-cell engager antibody. Cell viability was followed on the xCelligence device for 4 days, indicating that the redirected T-cells specifically killed all HBs-expressing Huh-7 cells. E Evaluation of IFN-γ in the supernatant of PBMCs co-cultured with HBs-expressing Huh-7 cells by ELISA.

To therapeutically target HBs on HBV-infected cells, T-cells can be redirected using a chimeric antigen receptor (CAR) (Bohne et al., 2008) or T-cell engager antibodies that contain the scFv C8 antibody targeting HBs and either a CD3 or a CD28 scFv activating T-cells (BiMAab) (Quitt, Luo et al, manuscript submitted for publication). Huh-7 cells transfected with plasmid expressing S, L or SML proteins served as target cells.

Human T-cells were engrafted with a scFv C8 based CAR (S-CAR) (Bohne, Chmielewski et al., 2008) by retroviral transduction. In line with earlier experiments (Krebs, Bottinger et al., 2013a), viability analysis revealed that engrafted T-cells specifically killed Huh-7 cells expressing the HBV S-protein or all three envelope proteins (SML) (Fig 6C). The same was observed when T-cells from healthy donors were redirected and specifically activated by the BiMab (Fig 6D). S-CAR T-cells as well as BiMab activated T-cells, however, also killed Huh-7 cells only expressing L protein preventing that T-cells can be activated by viral particles or SVP secreted (Fig 3C) and ensuring that only HBs located to the plasma membrane can be targeted (Fig 6C, D). Determining IFN-γ in cell culture supernatant showed that S-CAR T-cells were specifically activated in co-culture experiments (data not shown), but also T-cells were activated by T-cell engager BiMab upon binding to HBs on the target cell surface (Fig 6E). Both therapeutic approaches proved that HBV S, M and L are expressed on the surface of target cells and can be used to specifically target and kill HBs-expressing or HBV-replicating cells.

## Discussion

In this study, we could confirm the membranous distribution of HBs *in vivo* in HBV-transgenic mice and in liver and cancer tissue from chronic hepatitis B patients. Our data verified the membranous distribution in cell culture experiments revealing that HBs shows a speckled membrane-localisation in non-permeabilized, HBs expressing cells. Using a recombinant anti-HBs antibody, MoMab, we confirmed correct conformation of membrane-associated HBs, and intravenously injected MoMab localised to hepatocytes in the liver of Alb-Psx transgenic mice expressing HBs. By EM detecting membrane localisation of HBs in HBV-replicating hepatoma cells and in HBV-infected cells, we establish that membrane-associated HBs is derived from intracellularly expressed HBs, and does not result from the attachment of SVPs or virions. T-cells grafted with a CAR using the same scFv as the MoMab specifically eliminated Huh7 cells expressing HBV L-protein that is not secreted excluding the possibility that SVP or virions binding back to the cell membrane after secretion are targeted. Taken together, our study demonstrates that membrane-associated HBs enables the development of new and promising approaches to detect, monitor and treat HBV-infection.

In this study, we characterised the membrane-localisation of HBs *in vivo* staining hepatocytes from liver specimens of HBV-transgenic mice and chronically infected patients. Our results confirm a previous report performing indirect immunofluorescence on liver specimens. In tissue sections from 75 patients with chronic hepatitis B the authors found purely membranous or solely cytoplasmic as well as mixed staining of HBs on hepatocytes (Jiang et al., 2015). Moreover, this study indicated that the distinct membranous pattern of HBs is a sensitive and specific marker of active replication (Jiang et al., 2015). In another study, also patients with chronic hepatitis B including inactive carriers and cirrhosis patients accumulated HBsAg within the membrane of hepatocytes (Safaie et al., 2016b). It is well known that there is a correlation between hepatocarcinogenesis and high HBsAg levels in patients (Chisari, Klopchin et al., 1989, Sunami, Ringelhan et al., 2016). Dependent on the presence of HBeAg, HBs distribution seems to be different. The majority of the HBeAg-negative cases had a ‘patchy’ or membranous HBsAg distribution compared to only 50% among HBeAg-positive cases (Jiang et al., 2015, Lambert et al., 2007).

The punctate membrane-located HBs staining by IHC (Jiang et al., 2015, Lambert et al., 2007) needed to be verified in non-permeabilized hepatocytes with an intact plasma membrane. Therefore, an indirect immunofluorescence protocol with antibody staining before fixing the cells was established. By this approach, we confirmed a speckled membrane-distribution of HBs in HBV-replicating cells, but could neither detect this in non-infected HepG2 cells nor in HepG2 cells pre-incubated with HBV virions. The speckled HBs-localisation within the plasma membrane was confirmed by different HBs-specific antibodies such as HB01 and 5F9, which both recognise a linear epitope within the ‘a’ determinant of S as reported before (Berkower, Spadaccini et al., 2011, Inoue, Krueger et al., 2015). Using transiently transfected Huh7 cells expressing L but no S protein, we verified the punctate membrane-localisation of HBs independently of SVP secretion and circulating virions that maybe adsorbed (Chisari, Filippi et al., 1987, Roingeard & Sureau, 1998).

In previous studies we have shown that T-cells grafted with scFv C8 can specifically target cells infected with HBV or replicating HBV (Bohne et al., 2008, Krebs et al., 2013a). To visualise and demonstrate localisation of HBs to the plasma membrane, we constructed a new recombinant, human antibody that we named MoMab, containing two C8 binders attached to the constant region (CH2 and CH3) of human IgG1 based on this scFv. MoMab recognises a conformational epitope within the ‘a’ determinant of S verified using constitutive mutants of the AGL region and has the capability to neutralise hepatitis B virus infection. Staining with MoMab confirmed the speckled, membranous distribution of HBs on HBV-replicating and HBV-infected cells by immunofluorescence microscopy and by EM. Using S-CAR grafted T-cells, we demonstrated that not only cells expressing S or co-expressing all three envelope proteins are targeted but also cells that express only L protein. Furthermore, we found that intravenously injected MoMab could target hepatocytes in the liver of Alb-PSX mice expressing the HBV L protein. HBV L protein, when expressed alone, is not secreted (Chisari et al., 1987, Roingeard & Sureau, 1998). We demonstrate that L protein, when expressed in absence of the other HBs proteins, can still be targeted on the hepatocyte membrane of transfected cells and transgenic animals, indicating that the localisation of HBs on the membrane is independent of SVP or virion secretion. These data strongly support the hypothesis that HBs can be transported via MVBs that finally bud in the plasma membrane and transport the HBV envelope proteins to the cell membrane.

The endosomal sorting pathway via the formation of MVBs drives the release of infectious virions and filaments (Hoffmann et al., 2013, Jiang et al., 2015, Lambert et al., 2007, Stieler & Prange, 2014, Watanabe et al., 2007), whereas spheric SVPs are released by the constitutive secretion pathway (Huovila, Eder et al., 1992). Secretion of proteins via the endosomal sorting pathway is mediated by the endosomal sorting complex required for transport (ESCRT, consisting of ESCRT-0, ESCRT-I, ESCRT-II and ESCRT-III). It was described before, that L protein - like the Gag protein of HIV - is linked to the ESCRT-I component tsg 101 via alpha-taxilin (Hoffmann et al., 2013) and gamma 2-adaptin (Lambert et al., 2007) implying that L protein is involved in recruiting the ESCRT machinery and activating the ESCRT-MVB secretion pathway. These interactions could induce MVB-plasma membrane fission at the budding site, and thereby facilitate HBV release (Blumberg, 1977, Hoffmann et al., 2013, Huovila et al., 1992, Jiang et al., 2015, Lambert et al., 2007, Stieler & Prange, 2014, Watanabe et al., 2007). Interestingly, the assembly and release of SVPs derived from S alone occurs ESCRT-MVB-independent (Lambert et al., 2007). The mechanism of how HBs is involved in the ESCRT-MVB-driven exit of HBV is the major question of intense studies. It is still elusive, how HBs is incorporated in MVB, and thereby involved in the release of infectious virions and filaments from the infected hepatocyte.

In summary, we show that HBs - besides its intracellular localisation and secretion as spheric SVP – localises to the plasma membrane of HBV-infected cells where it is detected by antibodies recognising linear as well as confirmational epitopes, and can be targeted by redirected T-cells. Translocation to the plasma membrane most likely occurs via MVBs but further studies are required to validate this hypothesis.

## Materials and Methods

### Cell culture

The human hepatoma cell lines HuH7 (Nakabayashi et al., 1982) (JCRB0403), HepG2 (ATCC Cat# HB-8065), HepG2.2.15 (Sells, Chen et al., 1987) and HepG2-NTCP-K7 (Ko, Chakraborty et al., 2018) were grown in supplemented Dulbecco’s modified Eagle medium (DMEM) at 37°C in 5% carbon dioxide. DMEM medium was supplemented with 10% fetal calf serum (FCS), 2 mM L-glutamine, 1% NEM non-essential amino acids, 1% sodium pyruvate, 50 U/ml penicillin/streptomycin (all from Gibco, Thermo Fisher Scientific, Carlsbad, CA, USA). HBV-producing HepAD38 cells, kindly provided by Dr. Chris Seeger, were maintained in DMEM/F12 medium supplemented with 10% FCS, 1% penicillin/streptomycin/tetracycline and 0,4% G418 (all from Gibco, Thermo Fisher Scientific, Carlsbad, CA, USA) at 37°C in 5% carbon dioxide.

### Human biosamples and mouse lines used

Mice transgenic for LHBs (Alb-Psx) or a 1.3-fold overlength HBV genome (HBV1.3.3.2) were bred in a specific pathogen–free animal facilities and were bred and received humane care according to the German Law for the Protection of Animals with permission of the regulatory authority.

Human liver samples were obtained from liver parts directly after resection of hepatocellular carcinomas from patients chronically infected with HBV. They were banked in a professional biobank after fixation with 10% neutral-buffered formalin and paraffin embedding fulfilling Tier1 criteria of the BRISQ reporting guidelines (Biospecimen Reporting for Improved Study Quality). Samples were collected and processed with permission of the regulatory authority, authorized by the ethics committee of the University Hospital rechts der Isar of the Technical University of Munich (ref: 5846/13).

### Immunohistochemistry (IHC) staining

For HBs-staining, liver specimens of HBV-transgenic mice (strain HBV1.3.3.2) (Guidotti, Matzke et al., 1996), and chronic hepatitis patients that had developed hepatocellular carcinoma were fixed in 4% paraformaldehyde for 24 before paraffin embedding. 2 µm tissue sections were collected and stained by IHC. To analyse the location of HBs, sections were stained with monoclonal anti-HBs antibodies such as 70-HG15 (mouse) (Fitzgerald, Acton, MA, USA) and MoMab (human) and according secondary antibodies coupled to horseradish peroxidase (Thermo Fisher Scientific, Carlsbad, CA, USA). Histological analysis of the 2 µm paraffin-embedded tissue sections was performed with the Bond Polymer Refine Detection Kit (Leica, Wetzlar, Germany) on a BondMax system (Leica). Images of the HBs and MoMab staining’s were documented by scanning the whole slides using an SCN-400 slide canner (Leica) and analysed using Tissue IA image analysis software (Leica) with optimised quantification algorithms.

### Fluorescence analysis

To analyse the intracellular HBs distribution, transiently transfected Huh7 and HBV-positive HepG2.2.15 and Hep-AD38 cells were analysed by confocal microscopy. HuH7 or HepG2-NTCP cells were transfected with plasmids pSVB45H (expressing SML), pSVL (coding L only) or pSVBX24H (coding S) (Siegler & Bruss, 2013) or mCherry-S (expressing S fused to mCherry), kindly provided by Volker Bruss, using FuGENE® HD transfection reagent (Promega, Madison, WI, USA) for 48 hours (h) at 37°C. In addition, HBV-infected HepG2-NTCP and HepG2.2.15 cells were stained by indirect immunofluorescence to analyse the cellular location of HBs during viral replication. Parental HepG2 cells served as negative control. To detect HBs, cells were stained with anti-HBs antibodies HBVax (goat, dilution 1:500), HB01 (mouse, 1:200) (kindly provided by Prof. Dr. Glebe, University of Giessen, Germany), 5F9 (mouse, 1:200) (Golsaz-Shirazi, Mohammadi et al., 2016) and MoMab (human, 1:150) for 1h at 4°C and according secondary Alexa-Fluor® 647- or Alexa-Fluor® 568-conjugated antibodies (dilution 1:500) (Thermo Fisher Scientific, Carlsbad, CA, USA) for 45 min at 4°C. The samples were counterstained with the plasma membrane marker wheat germ agglutinin (WGA) coupled to Alexa-Fluor 488 (Thermo Fisher Scientific) diluted 1:500 for 30 min on ice. After washing the samples 3 times with 1% BSA in PBS, fixation was performed with 4% paraformaldehyde (PFA) for 10 min at room temperature (RT). For nuclear staining, coverslips were overlaid with DAPI Fluoromount-G® Mounting media (Southern Biotech, Birmingham, AL, USA). Images were documented by a confocal laser-scanning microscope (Fluoview FV10i, Olympus, Hamburg, Germany) and analysed using Image J (Schneider, Rasband et al., 2012) and Photoshop 7.0 (Adobe Software Palo Alto, CA, USA).

### Western blot analysis

HuH7 cells were transfected with plasmids pSVB45H, pSVL or pSVBX24H (Siegler & Bruss, 2013) using FuGENE® HD in OptiMEM™ reduced serum medium (Thermo Fisher Scientific, Carlsbad, CA, USA). Proteins from the whole cell lysates obtained after 24h, 48h and 72h were separated by SDS-PAGE and stained using the mouse monoclonal S-specific antibody HB01 (1:1000 dilution) overnight at 4°C as described before (Untergasser, Zedler et al., 2006). For loading control, the blot was incubated with a rabbit antiserum against ß-actin (1:10000 dilution) (42kDa, Sigma-Aldrich, Taufkirchen, Germany) for 1h at RT. Then, peroxidase-conjugated goat anti-mouse/-rabbit IgG antibodies (Sigma-Aldrich, Taufkirchen, Germany) were used at a dilution of 1:5000 and incubated for 2h at RT, followed by the enhanced chemiluminescence detection kit (Amersham Biosciences, Bath, UK).

### HBV infection and immunoassays

The supernatants of the transfected Huh7 cells were harvested after 24h, 48h and 72h at 37°C. HepG2-NTCP cells (seeded 5 × 10^4^ / 24-well plate) were differentiated with DMEM containing 1.8% DMSO for two days before infection with purified HBV at MOI of 200, 500 or 1000 virions by adding 4% polyethylenglycol over night as described (Ko et al., 2018). Media was exchanged every three days until the end of the experiment. At indicated time points post infection (p.i.), supernatants were collected and cleared by centrifugation at 5,000 g for 5 min, before HBeAg and HBs were determined on the Architect platform (Abbott, West Chicago, IL, USA). Total DNA and RNA were purified from infected cells using the NucleoSpin® Tissue and RNA kits, respectively (Macherey Nagel, Düren, Germany). RNA was transcribed into cDNA using SuperScript III reverse transcriptase (Invitrogen, Carlsbad, CA, USA). HBV DNA (rcDNA and cccDNA) and RNA (pregenomic RNA, pgRNA) were quantified by real-time PCR on a LightCycler™ instrument (Roche Diagnostics, Mannheim, Germany) using specific PCR primers as described (Lucifora, Xia et al., 2014, Untergasser et al., 2006).

### Purification and analysis of the human S-specific antibody MoMab

To produce the recombinant, anti-HBs antibody MoMab HEK 293 cells were transiently transfected and the antibody was purified by ion exchange chromatography from cell culture medium (InVivo Biotech Services, Berlin, Germany). Purified MoMab was separated by SDS-PAGE and stained with Coomassie Blue. For evaluation of the affinity constant (EC50), ELISA was performed using immobilised HBs overlaid with MoMab. For neutralization assay, HepG2-NTCP cells were infected as described above with HBV pre-incubated with MoMab (MOI of 500).

### Synthesis and physicochemical characterisation of MoMab-conjugated to super-paramagnetic iron oxide nanoparticles (SPIONs)

SPIONs were prepared from iron salt solutions by co-precipitation as described earlier (Shevtsov, Nikolaev et al., 2015). The surfaces of the obtained nanoparticles were functionalised with MoMab antibody producing superparamagnetic conjugates so-called MoMab-SPIONs. To analyse the size distribution and nanoparticle size of SPIONs and MoMab-SPIONs, transmission electron microscopy (EM) and dynamic light scattering (DLS) were performed. The hydrodynamic size and electrophoretic properties were measured on a Zetasizer Nano (Malvern Pananalytical, Malvern, UK). The NMR relaxation times of the *R*_2_*, *R*_2_ and *R*_1_ coefficients were detected by a CXP-300 NMR-spectrometer (Bruker, Billerica, MA, USA) with a magnetic field 7.1 Tesslar.

To assess the specific interaction of MoMab-SPIONs with HBs, the SPIONs and their obtained conjugates were co-incubated with recombinant HBsAg for 24h and the temporal T_2_ relaxation times was measured using the ‘switch assay’ as described (Tassa, Shaw et al., 2011).

### Transmission electron microscopy (EM)

HepG 2.2.15, HepAD38 cells and HBV-infected HepG2-NTCP cells (MOI of 100, experimental duration 9d) were trypsinized and seeded into a 48-well plate using 1×10^5^ cells/ml. After 3 d p.i., cells were incubated with SPIONs or MoMab-SPIONs for 3 h, washed 3 times with PBS and fixed using 2.5% glutaraldehyde (EM grade) in 0.1 M sodium cacodylate buffer pH 7.4 (Science Services, Munich, Germany). Samples were post-fixed in 2% aqueous osmium tetraoxide, dehydrated in gradual ethanol (30–100%) and propylene oxide followed by embedding in Epon (Merck, Darmstadt, Germany). Semithin sections were stained with toluidine blue, whereas the 50nm-ultrathin sections (on 200 mesh copper grids) were stained with uranyl acetate/lead citrate. Images were documented by EM (Zeiss Libra 120 Plus, Carl Zeiss NTS GmbH, Oberkochen, Germany) combined with a Slow Scan CCD camera and analysed using the iTEM software (Olympus Soft Imaging Solutions, Münster, Germany).

### S-CAR T-cell killing activity

HBV S-specific chimeric antigen receptor (S-CAR) was transduced into T-cells using retroviral vectors as described (Krebs, Böttinger et al., 2013b). HuH7 cells expressing SML, L or S after transient transfection were trypsinized, seeded into 96-well plates (3.5×10^3^ cells per well) and cultured overnight before S-CAR T-cells (1×10^6^ effector cells) were added. Cell viability was assessed over time for 4 days using the xCELLigence real-time cell analyser system and the RTCA software v1.2.1 (ACEA Biosciences, San Diego, CA, USA). To characterise T-cell activation, IFN-γ, TNF-α and IL-2 levels were quantified in co-culture supernatants by ELISA (BioLegend, San Diego, CA, USA).

### Animal experiments

Mice transgenic for LHBs (strain Alb-Psx; JAX stock # 027528) (Chisari, Filippi et al., 1986) or a 1.3-fold overlength HBV genome (strain HBV1.3.3.2) (Guidotti et al., 1996) were bred in a specific pathogen–free animal facilities receiving humane care. To evaluate, if MoMab specifically target HBs-expressing hepatocytes, 50µg and 100µg MoMab diluted in 100 µl PBS were intravenously injected into Alb-Psx mice or C57BL/6 mice (as control). 4 h after injection, mice were sacrificed and blood and liver were collected. Binding of MoMab to hepatocytes in liver specimens was examined by immunohistochemistry as described above. The study was conducted according to the German Law for the Protection of Animals with permission of the regulatory authority.

### Image and statistical analysis

Image analysis was performed using the Java-based image-processing program ImageJ 1.6.0 (NIH) (Schneider et al., 2012). The structural view of HBV S (PDB from van Hemert et al., 2008) (van Hemert et al., 2008) was generated using PyMOL molecular graphics system, version 1.3 (Schrödinger, LLC). The student *t* test statistical significance was calculated using GraphPad Prism, version 6.0c (GraphPad Software, San Diego, CA, USA).

## Data availability

This study includes no data deposited in external repositories.

## Acknowledgements

We thank Frank Thiele and Natalie Röder for their support with mouse experiments. We thank Romina Bester and Hortenzia Jacobi for HBeAg and HBsAg measurement, Chunkyu Ko, Shanshan Luo and Yuchen Xia for fruitful discussions, and Jochen Wettengel for providing HBV virus stocks. We are also grateful to Dr. Boris P. Nikolaev and Yaroslav Marchenko for NMR measurements.

## Author contributions

UP and BM designed the study and the experiments. LZ, FC, OQ, MF, MR, JH, FB and MA performed the experiments and acquired the data. LZ, FC, OQ, MF, CS, FB, MS and MK analyzed the data. CS, KW, VB, MS and FM provided experimental tools and interpreted the data obtained with these. LZ, MK, BM and UP drafted, wrote and revised the manuscript. All authors read and corrected the drafted manuscript and approved the submitted version.

## The paper explained

### Problem

Despite the availability of a prophylactic vaccine, Hepatitis B virus infection remains a global health problem causing more than 880,000 deaths each year. Once chronic, the infection cannot be cured by the available antiviral treatments that target the viral polymerase, and patients remain at risk to develop hepatocellular carcinoma. Therefore, new therapeutic strategies and alternative therapeutic targets are urgently needed.

### Results

Our study shows that HBV envelope proteins that primarily are embedded into the endosomal membrane of infected hepatocytes become translocated to the plasma membrane. They can be detected on the surface of HBV-infected and HBV-replicating cells, in liver sections of HBV-transgenic mice and HBV-induced liver cancer. A novel, recombinant antibody confirmed the correct confirmation of HBV envelope proteins on the cell surface and electron microscopy demonstrated their membrane distribution. Finally, we show that the envelope proteins can be recognised by redirected T-cells and allow targeting of infected hepatocytes by T-cell therapies using chimeric antigen receptors or T-cell engager antibodies.

### Impact

Our findings highlight that HBV envelope proteins are incorporated into the plasma membrane of infected hepatocytes and HBV-induced liver cancer cells where they provide an interesting target T-cell based, potentially curative therapies.

## Expanded View

**Figure EV1 - Intrahepatic distribution of HBs in HBV-infected patients**. Immunohistochemistry analysis of HBs distribution in biopsies of patients infected with HBV. In most samples we observed HBs was predominantly membrane-localised (A, white arrows), although in some samples HBs was mainly present in the cytoplasm (B, black arrows). Scale bars represent 50 μm.

**Figure EV2 - Characterisation of the conformational S-specific antibody MoMab**.

A Coomassie Blue staining of decreasing concentrations of purified MoMab separated under reducing (left panel) and non-reducing conditions (right panel).

B Alanine scanning of amino acids 101 to 225 of the S protein. Bars represent the interaction strength between the mutant and MoMab as determined by ELISA.

C HepG2 cells were transfected with a plasmid expressing a mCherry-S fusion protein and stained with MoMab before permeabilization. After permeabilization, cells were co-stained with ER-marker anti-Protein Disulphide Isomerase (PDI). The diffuse S-staining (red, mCherry autofluorescence) co-localised with the ER. MoMab only interacts with the S expressing cells.

D HepG2-NTCP cells were infected with HBV and stained for HBs using MoMab (green) before permeabilization and for the hepatitis B virus Core protein (red) after permeabilization. Scale bars represent 10 μm.

**Figure EV3 - Membrane-associated HBs correlates with infection rate**.

A HepG2-NTCP cells were infected with HBV at different MOIs and stained for HBs using MoMab (white, red in merged figures) before permeabilization. The cell membrane was co-stained with WGA (green) and nuclei were stained with DAPI (cyan). Scale bars represent 10 μm.

B, C Evaluation of the increasing infection efficiencies was performed using HBsAg and HBeAg at day 7 and 9 p.i. (B), as well as HBV marker at day 9 p.i. (C). The graphs represent the average and mean ± SD of one experiment performed in triplicate. S/CO: signal over cut-off.

**Figure EV4 - Characterisation of MoMab conjugated SPIONs**.

A Schematic overview of the MoMab-SPIONs nanoparticles and (B) the cluster-model induced by HBs.

C Measurement of T2 relaxation times using the ‘switch’ assay (temperature shift from 20°C (left panel) to 37°C (right panel) indicating the specific binding activity of MoMab-SPIONs to HBs.

**Figure EV5 - Transduction of S-CAR-grafted T-cells with and without interleukin IL2**. Real-time analysis of cell viability after exposure of S-CAR-grafted T-cells seeded onto Huh7 cells transfected with S, L and SML recorded by xCelligence.

